# Human intelligence relates to neural measures of cognitive map formation

**DOI:** 10.1101/2024.12.28.630614

**Authors:** R.M. Tenderra, S. Theves

## Abstract

Psychometric research on intelligence consistently identifies latent factors underlying performance correlations across cognitive tasks, with one general factor (g) explaining most variance and predicting general life outcomes. Their biological basis is yet unresolved, in particular with regard to the neural information processing mechanisms that may underlie intelligence. Here we test the hypothesis that interindividual differences in relational processing, supported by cognitive maps in the hippocampus, are related to fluid intelligence (g_f_), a statistical approximation of the general factor. Using standardized cognitive tests and fMRI of different mnemonic tasks, we demonstrate a positive correlation between g_f_ and map-like encoding of object positions learned piecemeal. The behavioral and neural geometry of object representations in the lower g_f_ range was less consistent with any two-dimensional representation, congruent with a lack of relational integration at encoding. The specificity of the link between cognitive maps and intelligence to relational processing was further supported by comparisons to non-relational mnemonic processing in the hippocampus during item recognition. These findings offer first empirical support for a link between neural mechanisms related to relational reasoning and general cognitive performance and probe the presumed relevance of hippocampal coding properties for cognition more broadly.

## Introduction

The study and definition of human intelligence is a longstanding interdisciplinary endeavor. Psychologists assess general cognitive ability by evaluating performance across diverse cognitive tasks^1^. A major discovery is that positive correlations between task performances can be explained by a small number of latent factors, which load onto one general factor g^2^. G is also the strongest single predictor of general life outcomes^3,4^. Among the second-order factors, the general ability for reasoning or fluid intelligence (g_f_) shows a near-perfect correlation to g^5^. While this factor structure is well-replicated, its biological basis remains unclear. Does a confined set of cognitive and neural functions underlie g? Previous neuroimaging research on intelligence has focused on locating interindividual differences in anatomical properties^6–8^, structural and functional connectivity^9,10^, or general activation levels, with an emphasis on the parieto-frontal cortices^11–13^. However, the mechanisms of neural information processing that may underlie general cognitive abilities are yet unknown. Here we take an information processing-based perspective, asking how individuals differ in the way they encode task information with respect to measures of general cognitive performance. Given the significant contribution of general reasoning ability to g^5,14^ and the central role of relational reasoning in human cognition^15–17^, it can be hypothesized that the general factor is related to the degree to which individuals encode the relational and structural information of experiences critical to ad-hoc inferences and transfer to novel situations^18^. Empirical and computational work implies a critical role for representational properties of the hippocampal-entorhinal system in reasoning: The system is thought to integrate spatial and non-spatial relations between experiences^19–25^ into domain-general cognitive maps of physical and abstract task spaces^26–28^, especially when these are task-relevant^20^. In particular, the hippocampal system may support decision making by compressing multivariate inputs according to task-relevant dimensions^29–31^. Such map-like, relational representations reflect relations between experiences as distance and direction between locations in physical or abstract task spaces on one or more latent dimensions and may thereby help capture relational structure from experiences in support of reasoning, i.e., infer unobserved links^32–34^ and states^35^ in the same context, or eventually generalize it to novel contexts. For instance, the hippocampus has been shown to support the inference of unobserved relations in paired associative inference^34,36^ and transitive inference tasks^32^ which rely on integrating individual associations into a collective representation of the underlying relational structure. Structural representations also allow alignment across problem instances, which can afford fast inferences via vector-based processing^36^. In this study, we set out to test whether interindividual differences in relational coding in the hippocampal system, associated with its function in forming and maintaining cognitive maps, predict fluid intelligence. To this end, we conducted comprehensive standardized cognitive tests as well as functional magnetic resonance imaging (fMRI) during different mnemonic tasks (including an established fMRI measure of cognitive map formation, the representation of two-dimensional distances between objects whose spatial location was learned piecemeal) in a large sample of participants and evaluated hippocampal processing with respect to intelligence factors. Our findings reveal a significant relation between map-like relational encoding of task information in the hippocampus and general reasoning ability g_f_.

## Results

### Performance in an object location memory paradigm

As a neural measure of map-like relational encoding, we assessed the hippocampal representation of two-dimensional (2D) distances between objects previously encountered in an object location memory task (Figure 1A-B). This measure is well established in both spatial and non-spatial tasks^22,29,37–39^, however, the spatial version of this paradigm offers distinct advantages, including moderate task difficulty and rapid learning speed, which help minimize dropout rates and maintain equivalent experimental procedures across a large participant sample.

**Figure 1.**
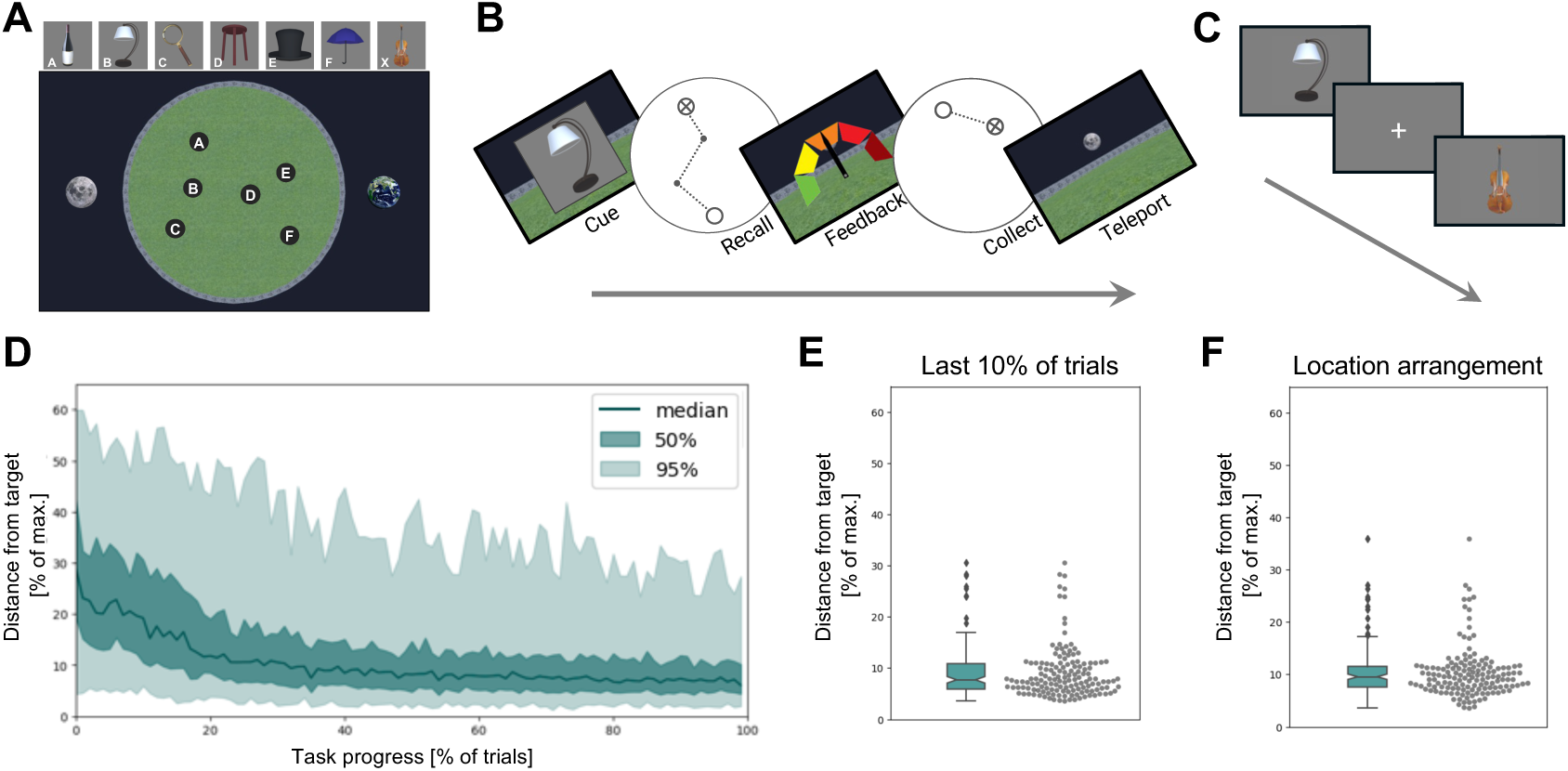
Performance in the object location memory paradigm. (A) Top-down view onto the virtual arena with diametrically opposed landmarks outside the arena. Each of the 6 locations A to F was associated with one object. Participants navigated the arena from a first-person perspective. (B) In each trial of the object location memory task, one of the objects had to be placed in the empty arena. Subsequently, feedback was displayed and the object appeared at its true location to be collected by the participant. (C) To measure object-specific neural response patterns all objects and an additional catch object were presented individually in the object picture viewing task. (D) Replacement errors during the object location memory task decreased and plateaued over the course of the task. The green line indicates the median error, shaded areas indicate the range covering 50% (dark green) and 95% (light green) of all data points. (E) Replacement error in the last 10% of trials was on average (SD) 9.1(5.04)% of the maximum possible distance (diameter). (F) Replacement errors in a post-learning object arrangement task, in which object locations had to be arranged from a top-down perspective, show a similar distribution.

In the object location memory task, participants learned to replace six cued objects in a circular arena. Each placement was followed by feedback and participants navigated to the true object location (Figure 1 A-B). Each trial began at a random starting position, leading participants to experience each object position independently, without navigating between objects. By the end of the learning phase, participants knew the positions of the objects well (performance in the last 10% of trials: mean(SD) = 9.36(5.26)% of max. distance; Figure 1D-E).

To assess hippocampal representations independent of spatial navigation demands, participants completed object viewing blocks before and after learning the associated object locations (Figure 1C). They detected the presence of a target object with high accuracy (percentage correct: mean(SD) = 98.6(2.7)%), affirming their attention to the stimuli. In a final assessment at the end of the experiment, participants recreated the object location arrangement from a top-down view of the arena. Their arranged locations closely matched the true locations (mean(SD) = 9.13(2.80)% of max. distance; Figure 1F), with inter-object distances between the arranged locations strongly correlating with the true inter-object distances (Pearson correlation: mean(SD) = 0.88(0.14)).

### Map-like relational encoding in the hippocampus scales with g_f_

We hypothesized that fluid intelligence (g_f_), the general ability for reasoning, relates to the extent to which individuals encode objects in a map-like, relational manner. Such relational map-like encoding would manifest in a neural representation of the 2D distances between objects, with higher multivoxel pattern similarity for closer objects (Figure 2A). To test this, we measured changes in hippocampal representations in object viewing blocks before and after location learning. Among other aspects of experimental control (see Methods), this design allows accounting for pre-existing object similarities unrelated to the information in the task. Consistent with group-level analyses in previous reports, we found a significant representation of the 2D distance relations between the objects. Hippocampal pattern similarities correlated with the 2D distance model matrix from the object location arrangement task (right hippocampus: t(54) = 2.47, p = 0.008; left hippocampus: t(54) = 0.57, p = 0.286; Figure 2B). As our main hypothesis posited a positive correlation with g_f_, the initial group-level test was performed on the upper third of g_f_-ranked participants. Consistent with our main hypothesis, we observed a significant positive correlation between the strength of the hippocampal 2D distance representation and g_f_ across participants. Higher g_f_ scores were associated with a stronger correspondence between hippocampal pattern similarity and the 2D distance model (right hippocampus: r(147) = 0.17, p = 0.021; Figure 2C). This correlation also persisted after accounting for behavioral performance in the object location memory, the location arrangement, and the picture viewing tasks (r_partial_(147) = 0.20, p = 0.008), indicating that variance in the neural representational format is not simply due to variance in the accuracy of available task information. Congruent with the main correlation of map-like representations to the factor g_f_, their correlation to individual cognitive tasks in the test battery scaled with the tasks’ respective g_f_-loading (r = 0.733, p < 0.001). Likewise, among all assessed cognitive abilities besides g_f_, an increased g_f_-loading leads to stronger correlations with map-like representations (r = 0.972, p < 0.001), supporting the notion that relational coding in the hippocampus is related to the latent construct g_f_ without being driven by a disproportionate relationship to any individual task. Figure 2D visualizes the neural representational space for the upper and lower thirds of the g_f_ distribution using Multidimensional Scaling (MDS), depicting lower congruency with the true location arrangement in the lower g_f_ group. Representations in the lower g_f_ spectrum may reflect noisier, yet map-like relational representations, or different ways of encoding could lead to systematic distortions. Here, we found no evidence in support of a map-like representation in the lower g_f_ group (t(57) = −0.88, p = 0.808).

**Figure 2.**
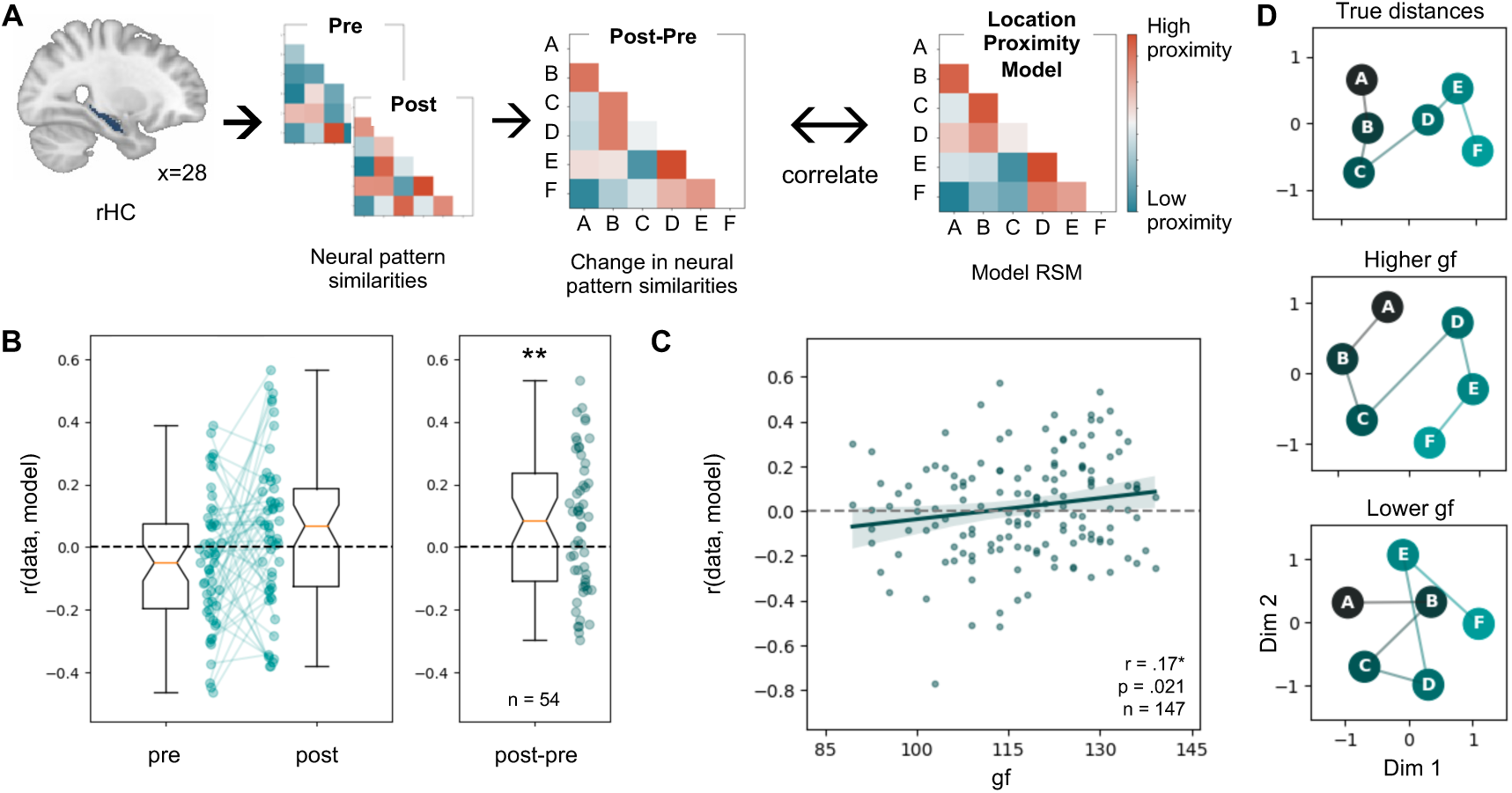
Map-like relational representation in the hippocampus scales with fluid intelligence (g_f_). (A) Measuring the representation of 2D distances between objects in the object location memory task. The baseline representational similarity matrix (RSM) acquired before the location learning phase was subtracted from the post-learning RSM to remove similarities unrelated to the information in the task. The resulting neural RSM was correlated with a location proximity model RSM. High correlations indicate that hippocampal response patterns to the objects became more similar for close locations and more different for far locations. (B) Significant 2D distance representation in the right hippocampus. (C) The hippocampal representation of the 2D arrangement of objects correlates positively with g_f_. The green line represents a linear regression fit; the green shadow represents the 95% confidence interval. (D) MDS visualizations of the distances between true locations (top) and the average neural pattern similarities for participants in the upper (middle) and lower (bottom) third of the sample’s g_f_ distribution.

### Geometrically inconsistent responses may indicate a lack of relational integration with lower g_f_

The reduced alignment of hippocampal representations with 2D object relations in the lower g_f_ group (Figure 2D) may particularly reflect a lack of relational integration at encoding, which can result in geometrically inconsistent beliefs about single object pairs that violate the constraints of 2D space. To explore this hypothesis, we first compared hippocampal representational dissimilarity matrices (RDMs) between the lower and higher g_f_ group, assessing the generic 2D-ness of the representational geometry, i.e., how well similarities between object-specific neural responses could be expressed as any 2D arrangement. To this end, we measured the variance explained by the first two Principal Components of a Principal Component Analysis (PCA). Consistent with our hypothesis, the first two PCs explained significantly more variance in the higher g_f_ compared to the lower g_f_ group (t(54,55) = 1.74, p = 0.042, Figure 3A). This effect was specific to the combined explained variance of the first two PCs, but did not reach significance for each PC individually (PC1: t(54,55) = 1.37, p = 0.087; PC2: t(54,55) = 0.80, p = 0.212). We further asked whether behavior was congruent with this explanation. In addition to the location arrangement task, participants also performed a pairwise distance estimation task (Figure 3B-C). This task is particularly informative in this context, as it queries object-to-object relations individually and, unlike the location arrangement task, does not require estimates to be overall geometrically consistent within the constraints of 2D space. Congruent with the neural analysis, also in the behavioral RDMs more variance was explained by the first two PCs in the higher g_f_ group compared to the lower g_f_ group (t(54,55) = 3.58, p < 0.001; cf. Figure 3D). While participants overall performed well in this task (mean(SD) = 0.70(0.20)), we further assured that the group difference in variance explained by the first two PCs is not based on inconsistent responses by replicating it after accounting for the coherence of responses per object pair (t(54,55) = 2.52, p = 0.007; cf. Figure 3E). Taken together, participants in the lower g_f_ group did not respond randomly but tended to show both behavioral and neural representations that were less consistent with a 2D geometry of the object relations than representations of participants in the higher g_f_ group. This indicates lapses in relational integration at encoding of individual locations, and possibly alternative memorization strategies, in the lower range of the g_f_ spectrum.

**Figure 3.**
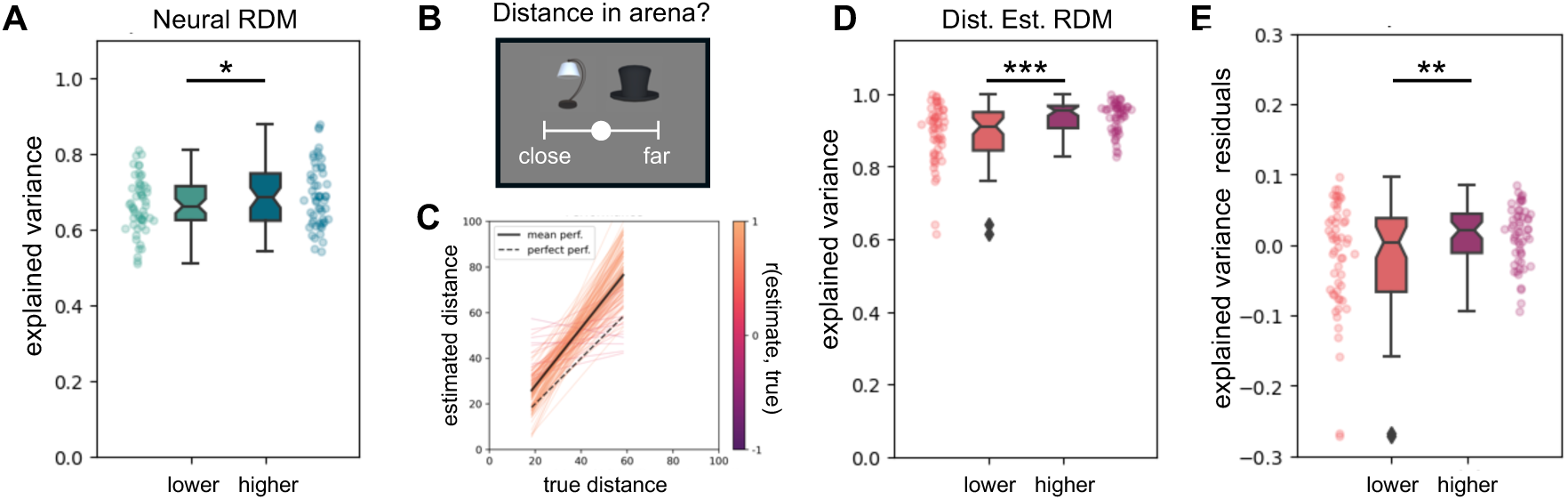
Neural and behavioral RDMs are less consistent with a 2D geometry in the lower g_f_ group. We assessed how much variance of the neural and behavioral representations of object-to-object relations is explained by the first two principal components (PCs) as an indicator of inconsistencies with a 2D integration of all relations. The more variance of the RDMs is explained by additional components, the less the represented pairwise distances reflect a 2D arrangement. (A) In the neural RDMs (Figure 2A), the first two PCs explain more variance in the higher g_f_ group than in the lower g_f_ group. (B) In a pairwise distance estimation task participants rated the distance between two presented objects using a slider from 0 (same location) to 100 (opposite ends of the arena). (C) The correlations between estimated distance and corresponding true distance are overall high (mean(SD) = 0.70(0.20)). (D) Congruent with (A), also in the behavioral RDMs of the pairwise distance estimation task, the first two PCs explain more variance in the higher g_f_ compared to the lower g_f_ group. (E) This is also the case after accounting for response coherence by comparing the groups on the residuals from regressing response coherence on explained variance by the first two PCs.

### The relation between hippocampal representations and intelligence is specific to relational processing

To further evaluate whether the relationship between intelligence and hippocampal processing is specific to the relational component of the representation, we replaced the neural measure with a non-relational mnemonic response in the same brain region (Figure 4B). To this end, participants performed an item recognition memory task during fMRI in which they indicated whether objects had been presented in a preceding encoding task (Figure 4A). As established in previous studies^40–42^, we observed a significantly higher BOLD response in the right posterior hippocampus for correctly recognized compared to correctly rejected novel items (t(154) = 4.39, p < 0.001; Figure 4B). This non-relational item memory signal did not significantly correlate with g_f_ (r(154) = 0.07, p = 0.183), and this correlation was significantly lower than the correlation between g_f_ and relational map-like representations (z(130) = 1.689, p = 0.047).

**Figure 4.**
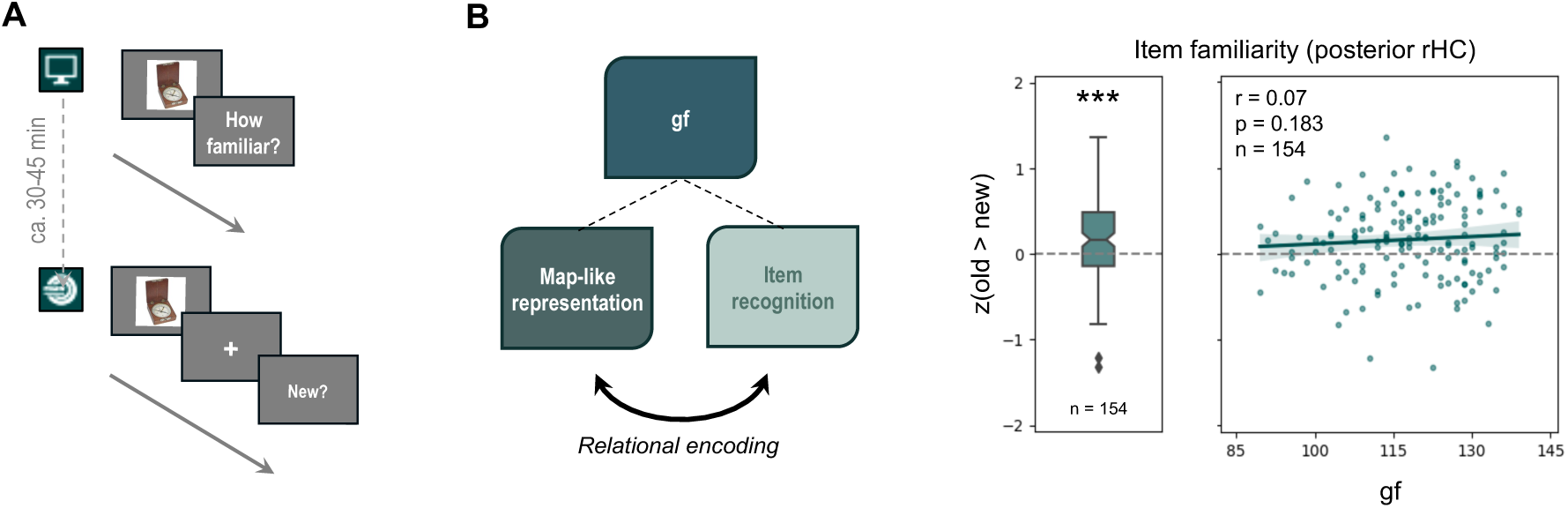
The relation between intelligence and cognitive maps is specific to relational processing. (A) As a non-relational mnemonic signal in the hippocampus, we measured the item recognition effect in a paradigm, in which participants indicated whether an item was old or new with respect to a preceding implicit encoding phase. (B) To test whether the relationship between g_f_ and neural representations in the hippocampus is functionally specific to relational processing, we compared it to the relation between g_f_ and non-relational item memory in the hippocampus. The non-relational item recognition memory signal in the right hippocampus did not correlate with g_f_.

## Discussion

How do individuals differ in the way they process information with respect to their general cognitive ability? We set out to examine general factors underlying multi-task performance, in particular general reasoning ability (g_f_), for the first time, on the level of neural representational properties. Our findings reveal that hippocampal cognitive maps, a representational format integrating relations along multiple dimensions, correlate with g_f_. We further show that this relation is specific to the relational component of processing and g_f_, the factor with the highest g-loading among all broad abilities^5,14^.

We use the representation of 2D distances between locations in physical or abstract spaces as an established measure of map-like relational representations^22,29,37–39^. We found that individual model fits positively correlated with the factor scores for general reasoning ability (g_f_), as measured via a standardized test battery, suggesting that relational encoding was more pronounced in individuals with higher g_f_ scores. Notably, this correlation persisted even when accounting for interindividual differences in task performance (i.e., in the object location memory and object viewing task), indicating that variance in the neural representational format is not merely due to variance in the availability of task information (i.e., how well locations were known or if object cues were attended). If relational coding in the hippocampus can be considered a g_f_/g-related mechanism, it could in principle also correlate with task performance. However, the influence of interindividual processing differences on task performance can depend on task difficulty. Here, we designed the task to be sufficiently easy for most participants to accurately learn the object locations within the given time and decouple individual differences in neural coding strategies from the availability of task information. Crucially, relational representations in the hippocampus were not enforced by task demands: Each object was encountered in isolation within the arena, with no explicit relationships between objects experienced. While participants could have integrated object relations into a unified map-like representation, they also could have relied on alternative navigation and memorization strategies (e.g., associating locations with landmarks or route-following from a certain reference point). Thus, the observed differences in neural representations rather reflect individuals’ tendencies to integrate information into relational representations.

Relatedly, interindividual differences in navigation strategies have been associated with both spatial and verbal working memory^43^, further supporting the view that cognitive map-based navigation improves spatial learning and navigation performance^44^ and links to higher order cognitive abilities^45^. Beyond this spatial domain, tasks probing associative memory or relational integration at the behavioral level, have been shown to predict fluid intelligence above and beyond working memory^46,47^.

While we focus on the neural encoding of newly acquired information, differences in the encoding of knowledge may also affect the organization of consolidated semantic information in the long term, possibly reflecting in systematic g_f_-related differences observed in the relational structure of semantic networks^48^.

To better characterize the interindividual processing differences along the g_f_ spectrum, we compared the representational geometry of two extreme groups corresponding to the upper and lower third of the g_f_ distribution. If the relation between map-like representations in the hippocampus and g_f_ specifically reflects an effect of relational processing, lower g_f_ values should be associated with lapses in relational integration of separately encoded object locations. Such lapses could lead to representations of individual relations that violate the constraints of a 2D geometry. Consistent with this hypothesis, both neural and behavioral representations of object relations were less well explained by the first two principal components in a PCA in the lower g_f_ group compared to the higher g_f_ group. Together, these findings provide evidence for a link between neural coding of relational structure and g_f_.

Further analysis confirmed that the correlation between intelligence and hippocampal representations was specific to relational integration, as no such relation was observed for non-relational item recognition signals in the hippocampus. In sum, our results support the idea that neural mechanisms related to relational reasoning contribute to human intelligence.

We focused on the hippocampal-entorhinal system due to its central role in relational processing, organizing spatial and non-spatial relations into domain-general cognitive maps^27,49^ affording inference and generalization. In particular, computational modeling suggests that firing properties of entorhinal cells such as grid cells reflect abstract, specifics-independent coding of relational structure, which allows its flexible conjunction with the specifics of new situations by hippocampal place cells^28^. It was further observed that the hippocampus can represent task variables in a disentangled geometry, which is optimal for cross-context generalization^50^ and can emerge immediately after instruction^51^ and swiftly adapt to new goals or contexts^52^. Further investigations could consolidate the present findings by examining structural generalization in the human hippocampus and other regions supporting complementary functions with respect to intelligence.

The hippocampal-entorhinal system serves as a multimodal convergence hub^53–55^, well positioned to interact with domain-specific regions to constitute a general processing factor across diverse tasks.

Prior neuroimaging research on intelligence has largely highlighted relations of intelligence to the anatomical properties and overall activation levels in parietal and frontal cortices^11–13^. Interestingly, parieto-frontal regions are functionally coupled with the hippocampus and have shown similar^56,57^ as well as complementary^58^ representations in spatial and nonspatial tasks^59,60^, although their specific role requires further clarification.

Our results suggest a specific link between map-like relational representations of task information in the hippocampus and interindividual differences in general reasoning ability, an approximation of the general factor of intelligence. This suggests that the general factor may relate to the extent to which individuals tend to extract relational structure from experiences, which is a basis for flexible reasoning^61^. More generally, our approach showcases how psychometric insights about cognitive factors can help to probe the general relevance of neural processing mechanisms - here hippocampal representations subsumed under the framework of cognitive maps - for cognition.

## Acknowledgments

S.T. is supported by a Minerva Fast Track Fellowship of the Max Planck Society. R.T. is enrolled in the Max Planck School of Cognition. We thank all research assistants and technical support staff who supported the data acquisition. We thank Alexander Eperon and Max Hinrichs for sharing code, and Volker Reisner, Misun Kim, and Kim-Laura Speck for technical help on the VR task.

## Author Contributions

S.T. conceived the study. R.T. and S.T. designed and implemented the experimental tasks. R.T. lead the data acquisition. R.T. and S.T. analyzed the data. R.T. and S.T. jointly wrote the manuscript.

## Methods

### EXPERIMENTAL MODEL AND STUDY PARTICIPANT DETAILS

We recruited 179 healthy adults (90 female, 86 male, 3 diverse; mean age(SD) = 26.49(4.85)) who were right-handed, had normal or corrected-to-normal sight, spoke German natively or learned it before age 7, and held a high school diploma or equivalent. Participants provided written informed consent before participation. This study was approved by the ethics commission of the Medical Faculty of the University of Leipzig and adhered to the Declaration of Helsinki. The necessary sample size was approximated a priori based on an analysis of statistical power for correlations (r = 0.2, β = 0.8, p = 0.05).

### METHOD DETAILS

#### Experimental Procedure

Participants completed standardized cognitive test batteries to assess intelligence factor scores on a first day and participated in computerized tasks and MRI recordings on two other days (see below for a detailed description of the tasks).

#### Cognitive Tests

Participants completed all modules of the Intelligence-Structure-Test (I-S-T 2000 R), a German, standardized intelligence test battery^62^, measuring fluid intelligence (g_f_) and crystallized intelligence (g_c_) across verbal, numerical, and figural domains, as well as short-term memory (cf. Table S1 for individual task descriptions). A g_f_ score was determined according to the manual’s instructions by transforming each task score to reflect the participant’s performance relative to a reference population (a representative sample of adults from the German general population) weighted by their respective g_f_ loading, which were then entered into a sum to determine g_f_. Additionally, the test allows for the measurement of domain-specific cognitive abilities with lower g loadings (cf. Table S2 for factor descriptions). Processing speed was assessed via the Zahlen-Verbindungs-Test (ZVT), a standardized trail-making test ^63^.

#### Object Location Memory Task

The navigation tasks using the circular arena were built using Unity Game Engine (version 2018.1.9f2). All other tasks were implemented using PsychoPy (version 2021.1.3).

For all spatial navigation tasks, participants controlled a virtual character in a circular arena from a first-person perspective on a screen. Two distal landmarks were projected at infinite distance, and a boundary along the arena perimeter served as orientation cues. During familiarization, participants collected traffic cones until they felt comfortable handling the buttons. Subsequently, they learned 6 object locations. The object-location associations were randomized across participants. Participants collected each of the objects once at their true location before the object location memory task began. In each trial, participants were presented with a target object, which they replaced as close as possible to its true location in the empty arena. Participants received feedback about their replacement error (distance to the true target location). Then the object appeared at its true location, where participants collected it to start the next trial. After collection, participants’ characters were teleported to a new start location. Participants completed 3 learning task runs (up to 72 trials/run; 20.5 min/run). To illustrate the median learning curve, we applied linear interpolation to resample the varying number of drop errors for each participant (n_mean_(SD) = 177.0(28.5)) into a common range of 50 samples each representing a 2% increment in task progress. For each step, we computed the median value as well as the percentiles including 50% and 95% of all data points to capture central and broader distribution trends. To assess learning outcomes, we calculated the average drop error of all trials across the final 10% of trials for each participant (n_mean_(SD) = 18.0(2.8)).

#### Object Picture Viewing Task

We recorded fMRI data during a pre- and post-learning object picture viewing task from 179 participants. The pre- and post-learning runs used identical stimulus sequences. Participants viewed 6 objects previously encountered in the object location memory task along with one additional catch object. To probe attention, participants were instructed to press a button whenever a catch object appeared. Each object image was presented for 1.5 sec, followed by a fixation cross for an average duration of 4 sec (onset jittered using a truncated exponential function). All 7×7 possible image-to-image transitions occurred once before repeating, resulting in 196 trials including 14.14% catch trials. Each run lasted approximately 18 min. Participants were excluded from fMRI data analysis if they exhibited excessive motion (mean framewise displacement > 0.4), responded incorrectly in more than 75% of catch trials in any run, or had an average response time slower than 1.5 s in any run, as these criteria indicated insufficient attention to the stimuli.

#### Pairwise Object Distance Estimation Task

Following the Object Picture Viewing Task, participants completed behavioral post-learning tasks to assess their knowledge of the object locations in greater detail. In the distance estimation task, each possible combination of object pairs was presented twice, with object positions (left/right) counterbalanced across trials, in a pseudorandom sequence. Participants estimated the distance between objects on a scale ranging from 0 (indicating the objects share the same location in the arena) to 100 (indicating the objects were located at opposite ends of the arena’s diagonal). To identify participants who may have responded randomly, we simulated 100,000 datasets with object distance estimates drawn from a uniform random distribution. 2 participants were excluded because their response coherence (mean absolute difference between same-pair distance estimates) did not significantly differ from the distribution of random values, i.e. their coherence was not lower than the 99^th^ percentile of the random distribution. Performance in the distance estimation task is measured as the Pearson correlation between participants’ estimated distances and corresponding true distances.

#### Object Location Arrangement Task

Finally, participants completed an object location arrangement task which provided a top-down view onto the empty arena and landmarks. In Phase 1, participants arranged 6 grey circles to reflect the object positions. The circles initially appeared at the arena’s center and could be moved via drag and drop. In Phase 2, the locations were highlighted one by one and participants assigned an object to each location from a dropdown menu listing all 6 objects. To ensure meaningful object location arrangements, we excluded participants if their arrangement did not reflect 6 unique locations. Specifically, 4 participants did not complete the task as instructed failing to remove 2 or more circles from the arena center, and 9 participants assigned the same object to multiple locations. Performance in the object location arrangement task was determined by calculating for each participant the mean of the absolute distance between each participant’s arranged locations to the corresponding true locations.

#### Item Recognition Task

We collected fMRI data from 173 participants during an item recognition memory task. Note that due to technical issues, procedural errors, or participant dropouts, not all participants contributed usable data for both fMRI tasks. Participants were initially familiarized with a set of 180 unique everyday items ^64^ under the guise of a cover task. They were instructed to rate how often they had encountered each item or similar images in everyday life as quickly and intuitively as possible. Each item was displayed for 1 second, followed by the rating question. Approximately 30–45 minutes later, participants completed the item recognition memory task while undergoing fMRI scanning. In this task, 60 previously seen (“old”) and 60 novel (“new”) items were presented in a randomized sequence lasting approximately 13 min. Each item appeared for 1 sec, followed by a fixation cross for an average of 0.5 sec, and then a question mark prompting a response within 2 seconds. Participants indicated their recognition response (“yes” or “no”) and their confidence (“certain” or “uncertain”) via button press. An inter-trial interval followed, during which a fixation cross was displayed for an average of 3 seconds (jittered using a truncated exponential function). The same fMRI exclusion criteria were applied as in the object picture viewing task.

#### MRI

In the MRI scanner tasks were presented using a projector on a screen outside the scanner, which participants could view through a mirror attached to the head coil. Participants responded using MRI-compatible button boxes. MRI data was recorded using a 3T Siemens MAGNETOM Skyra Connectom A (Siemens, Erlangen, Germany) using a 32-channel head coil.

##### Anatomical imaging sequence

High-resolution T1-weighted anatomical images were acquired using a Magnetization Prepared 2 Rapid Acquisition Gradient Echo (MP2RAGE) sequence. The following acquisition parameters were used: repetition time (TR) = 5 s, echo time (TE) = 2.84 ms, inversion times (TI1, TI2) = 700 ms and 2500 ms, and flip angles = 4° (first readout) and 5° (second readout). The field of view (FOV) was 256 × 240 mm² with 176 sagittal slices, providing an isotropic voxel resolution of 1 mm³. Full k-space sampling was performed with a partial Fourier factor of 1. To produce the final image the INV1 and INV2 images were combined, then the final T1-weighted image was generated by multiplying the intensity-normalized INV2 image with the MP2RAGE UNI image using an AFNI-based script (3dMPRAGEise, https://github.com/srikash/3dMPRAGEise). This approach enhances tissue contrast while suppressing background noise and residual artifacts.

##### Functional imaging sequence

All fMRI data was recorded using a CMRR multiband echo-planar-imaging (EPI) pulse sequence (TR = 1.5s, TE = 24ms, flip angle = 70°, partial Fourier = 0.75, multiband-factor = 4, phase encoding direction = AP, voxel size = 2 mm^3^ isotropic, FOV = 204×204 mm^2^, 80 slices, interleaved slice acquisition order). The measurement volumes were angled parallel to the axis crossing the anterior and posterior commissure (ACPC). For each fMRI recording of a task, a corresponding EPI field map for estimation of the B0 field inhomogeneities was constructed based on 2 EPI reference images with opposite phase encoding direction.

#### Preprocessing

Raw DICOM files were converted into BIDS format using the dcm2bids package (version 2.1.7, https://github.com/UNFmontreal/Dcm2Bids). Results included in this manuscript come from preprocessing performed using *fMRIPrep* 22.0.1 (Esteban, Markiewicz, et al. (2018); Esteban, Blair, et al. (2018))

##### Preprocessing of B_0_ inhomogeneity mappings

A *B_0_*-nonuniformity map (or *fieldmap*) was estimated based on two (or more) echo-planar imaging (EPI) references with topup (Andersson, Skare, and Ashburner (2003); FSL 6.0.5.1:57b01774).

##### Anatomical data preprocessing

The T1-weighted (T1w) image was corrected for intensity non-uniformity (INU) with N4BiasFieldCorrection (Tustison et al. 2010), distributed with ANTs 2.3.3 (Avants et al. 2008, RRID:SCR_004757), and used as T1w-reference throughout the workflow. The T1w-reference was then skull-stripped with a *Nipype* implementation of the antsBrainExtraction.sh workflow (from ANTs), using OASIS30ANTs as target template. Brain tissue segmentation of cerebrospinal fluid (CSF), white-matter (WM) and gray-matter (GM) was performed on the brain-extracted T1w using fast (FSL 6.0.5.1:57b01774, RRID:SCR_002823, Zhang, Brady, and Smith 2001). Volume-based spatial normalization to a standard space (MNI152NLin6Asym) was performed through nonlinear registration with antsRegistration (ANTs 2.3.3), using brain-extracted versions of both T1w reference and the T1w template. The following template was selected for spatial normalization: *FSL’s MNI ICBM 152 non-linear 6th Generation Asymmetric Average Brain Stereotaxic Registration Model* [Evans et al. (2012), RRID:SCR_002823; TemplateFlow ID: MNI152NLin6Asym], *ICBM 152 Nonlinear Asymmetrical template version 2009c* [Fonov et al. (2009), RRID:SCR_008796; TemplateFlow ID: MNI152NLin2009cAsym].

##### Functional data preprocessing

For each of the 3 BOLD runs found per subject (across all tasks and sessions), the following preprocessing was performed. First, a reference volume and its skull-stripped version were generated using a custom methodology of *fMRIPrep*. Head-motion parameters with respect to the BOLD reference (transformation matrices, and six corresponding rotation and translation parameters) are estimated before any spatiotemporal filtering using mcflirt (FSL 6.0.5.1:57b01774, Jenkinson et al. 2002). The estimated *fieldmap* was then aligned with rigid-registration to the target EPI (echo-planar imaging) reference run. The field coefficients were mapped on to the reference EPI using the transform. BOLD runs were slice-time corrected to 0.702s (0.5 of slice acquisition range 0s-1.41s) using 3dTshift from AFNI (Cox and Hyde 1997, RRID:SCR_005927). The BOLD reference was then co-registered to the T1w reference using mri_coreg (FreeSurfer) followed by flirt (FSL 6.0.5.1:57b01774, Jenkinson and Smith 2001) with the boundary-based registration (Greve and Fischl 2009) cost-function. Co-registration was configured with six degrees of freedom. Several confounding time-series were calculated based on the *preprocessed BOLD*: framewise displacement (FD), DVARS and three region-wise global signals. FD was computed using two formulations following Power (absolute sum of relative motions, Power et al. (2014)) and Jenkinson (relative root mean square displacement between affines, Jenkinson et al. (2002)). FD and DVARS are calculated for each functional run, both using their implementations in *Nipype* (following the definitions by Power et al. 2014). The three global signals are extracted within the CSF, the WM, and the whole-brain masks. The head-motion estimates calculated in the correction step were also placed within the corresponding confounds file. The confound time series derived from head motion estimates and global signals were expanded with the inclusion of temporal derivatives and quadratic terms for each (Satterthwaite et al. 2013). Frames that exceeded a threshold of 0.5 mm FD or 1.5 standardized DVARS were annotated as motion outliers. Additional nuisance timeseries are calculated by means of principal components analysis of the signal found within a thin band (*crown*) of voxels around the edge of the brain, as proposed by (Patriat, Reynolds, and Birn 2017). The BOLD time-series were resampled into standard space, generating a *preprocessed BOLD run in MNI152NLin6Asym space*. First, a reference volume and its skull-stripped version were generated using a custom methodology of *fMRIPrep*. All resamplings can be performed with *a single interpolation step* by composing all the pertinent transformations (i.e. head-motion transform matrices, susceptibility distortion correction when available, and co-registrations to anatomical and output spaces). Gridded (volumetric) resamplings were performed using antsApplyTransforms (ANTs), configured with Lanczos interpolation to minimize the smoothing effects of other kernels (Lanczos 1964). Non-gridded (surface) resamplings were performed using mri_vol2surf (FreeSurfer).

Many internal operations of *fMRIPrep* use *Nilearn* 0.9.1 (Abraham et al. 2014, RRID:SCR_001362), mostly within the functional processing workflow. For more details of the pipeline, see [the section corresponding to workflows in fMRIPrep’s documentation](https://fmriprep.org/en/latest/workflows.html).

#### Regions of Interest

Regions of interest (ROI) were created based on a probabilistic map of HC from the Havard-Oxford Atlas (Makris et al. 2006, Frazier et al. 2005, Desian et al. 2006, Goldstein et al. 2007). Probability maps were thresholded at 80%. Anterior and posterior ROIs were created by dividing each HC ROI along the anterior-to-posterior axis of MNI space such that the resulting ROIs contained approximately equal numbers of voxels. Analyses of HC were performed on separate hemispheres, due to a commonly discussed lateralization of functions^65^.

#### First-Level Modelling of fMRI Data using General Linear Models (GLMs)

fMRI data analysis was performed in MNI space. A separate GLM was constructed for each task. For each object picture viewing run the model specified one regressor per object, whereas for the item recognition memory run four regressors were included to capture the conditions of correctly and incorrectly classified *old* and *new* items. Both GLMs incorporated the same set of nuisance regressors to account for non-task-related variability. These included regressors for button presses, six motion parameters derived from realignment, and cosine drift regressors to implement a high-pass filter with a cutoff frequency of 0.01 Hz.

### QUANTIFICATION AND STATISTICAL ANALYSIS

All hypotheses were directional and corresponding test statistics were based on one-sided tests. When analyses were performed on both hemispheres, we corrected the alpha-level accordingly.

#### Representational Similarity Analysis (RSA)

Representational Similarity Analysis (RSA) was used to investigate the neural representation of 2D spatial relations between the objects in the task. Object-specific response patterns were extracted from the first-level GLM for both the pre- and post-learning runs. These patterns were used to construct Representational Similarity Matrices (RSMs) for each run. RSMs were computed by calculating the Pearson correlation of neural response patterns for each object pair. The pre-learning RSM was subtracted from the post-learning RSM for each participant to remove pre-existing similarities and capture the learning-related representational changes. Each participant’s RSM was then correlated with a model RSM using Spearman’s rho. The model RSM entailed the estimated distances from the location arrangement task, accounting for minor variance in the perception of the arena (e.g., participants reporting not perceiving it as perfectly circular). Measuring object-location representations in this design allows separating location learning from the probing of location representations and it accommodates interindividual differences in learning speed and duration without affecting the neural measure itself. Unlike the learning task, the object viewing blocks ensure that all participants complete a consistent number of trials, and further reduce potential confounds from concurrent hippocampal processing related to navigation or learning.

#### Multi-Dimensional Scaling (MDS)

To visualize the representational spaces in individuals with low and high g_f_ scores, we split the data into thirds (corresponding to a g_f_ equal to or greater than the 67.7^th^ percentile in the higher g_f_ group, and a g_f_ equal to or lower than the 33.3^rd^ percentile in the lower g_f_ group). We used metric MDS to illustrate a 2D embedding that corresponds well to the multidimensional neural distance representations of the higher and lower g_f_ groups.

#### Dimensionality Analysis

To investigate whether neural RDMs in the low relative to the high g_f_ group reflect inconsistencies of relations with the constraints of a 2D arrangement, we assessed the variance explained by the first two principal components of a Principal Component Analysis (PCA) of each subject’s neural RDM. Any 2D arrangement of the object set should result in an RDM in which 100% of the variance is explained by the first two PCs. A larger proportion of variance explained by additional components indicates that the inter-object distances are less consistent with a 2D structure. Further, we conducted this analysis on the behavioral RDMs of participants’ pairwise object distance estimations. Repeated estimates per pair were averaged to construct the RDM. The effect of response coherence on the explained variance by the first two PCs was partialled out using a linear model predicting explained variance with response coherence and repeating the analysis on the residuals.

#### Item Recognition Memory

To quantify item familiarity responses in the hippocampus, we contrasted correctly classified old items and correctly classified new items (old_correct_ > new_correct_) and averaged the corresponding z-scores of the ROI for each participant.

#### Correlation of Neural Response Measures with g Factors

For correlation analyses between neural response measures and g factors, we computed Pearson correlations.

To compare correlations, we calculated Steiger’s z, used for comparing dependent correlations within the same sample. When comparing correlations from overlapping, but not identical samples—such as g_f_ with relational representations from the object location memory task versus non-relational representations from the item recognition memory task—we subsampled participants who completed both tasks to meet the test’s assumptions.

## SUPPLEMENTARY

S1. Decomposition of the Intelligence Test Battery

**Table S1:**
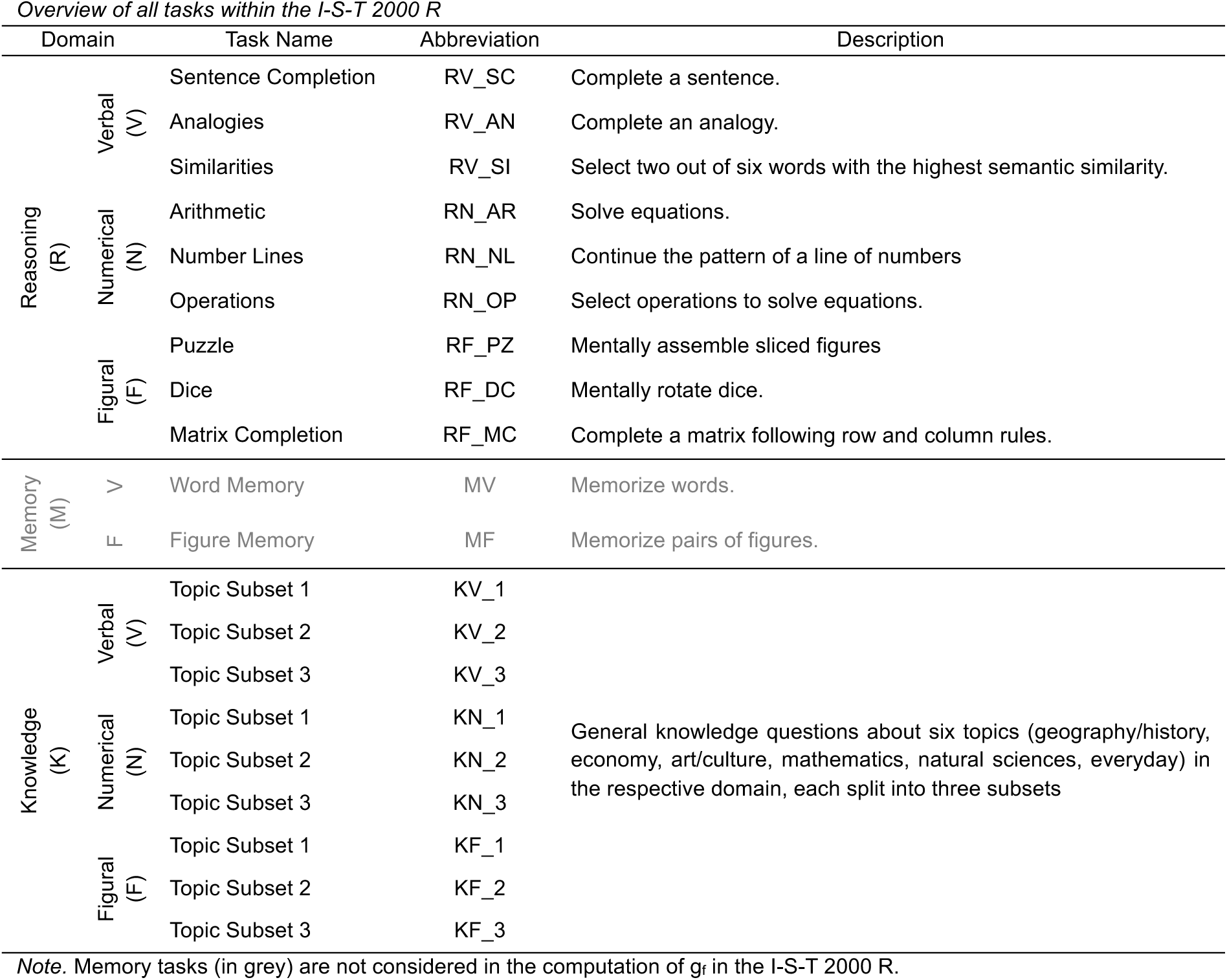
Overview of all tasks within the I-S-T 2000 R.

**Table S2:** Overview of measured cognitive abilities within the I-S-T 2000 R.

| Factor Name | Abbreviation | Description |
| --- | --- | --- |
| Verbal Intelligence |  | The ability to handle verbal material during inferential reasoning. The level of language fluency (i.e., vocabulary) and the ability to infer relationships between words both contribute to this ability. |
| Numerical Intelligence |  | The ability to solve arithmetic problems and infer logical relationships between numerical values. |
| Figural Intelligence |  | The ability to handle figural or spatial information in two- or three-dimensional space. This includes the ability to comprehend proportions of planes and volumes, as well as the ability to infer logical relations between figures. |
| Memory | $g_{\text{memory}}$ | The ability to actively encode information and retrieve it over a short period. |
| Crystallized Intelligence | $g_c$ | The accumulated knowledge deemed culturally relevant (broadly within Germany) |
| Fluid Intelligence | $g_f$ | The ability to infer relationships between pieces of information and engage in formal-logical, inductive, and deductive reasoning. |

**Figure S1.**
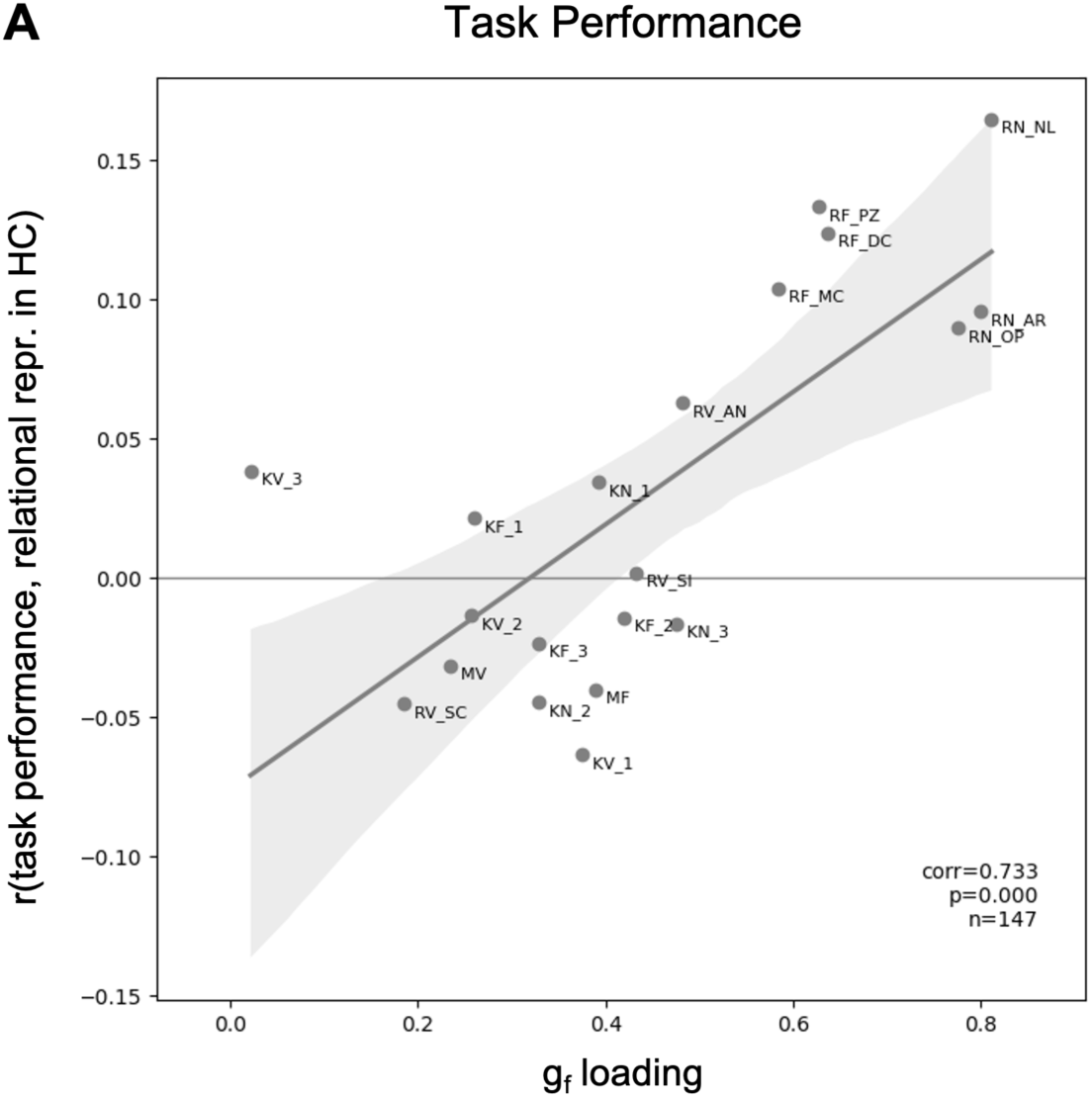
Decomposition of the test battery reveals a relationship between gf loading and relational representations. (A) All tasks of the I-S-T 2000 R (reasoning (R), memory (M), and knowledge (K) tasks, from the verbal (V), numerical (N), and figural (F) domains) are arranged based on their g-loading and their correlation with relational representations in Hippocampus (HC). A full description of task abbreviations is included in the supplementary material of the revised manuscript. With increasing gf loading of tasks, the respective task performance score correlates stronger with relational representations in HC. Abbreviations: RV_SC: Sentence Completion; RV_AN: Analogies; RV_SI: Similarities; RN_AR: Arithmetic; RN_NL: Number Line; RN_OP: Operations; RF_PZ: Puzzle; RF_DC: Dice; RF_MC: Matrix Completion; MV: Verbal Memory; MF: Figural Memory; KV_1-3: Verbal Knowledge; KN_1-3: Numerical Knowledge; KF_1-3: Figural Knowledge (cf. Table 1 for task descriptions).

